# Functional imaging of microbial interactions with tree roots using a microfluidics setup

**DOI:** 10.1101/506774

**Authors:** Marie-Francoise Noirot-Gros, Shalaka V. Shinde, Chase Akins, Jessica L. Johnson, Sarah Zerbs, Rosemarie Wilton, Ken Kemner, Philippe Noirot, Gyorgy Babnigg

## Abstract

Coupling microfludics with microscopy has emerged as a powerful approach to study at cellular resolution the dynamics in plant physiology and root-microbe interactions. Most devices have been designed to study the model plant *Arabidopsis thaliana* at higher throughput than conventional methods. However, there is a need for microfluidic devices which enable *in vivo* studies of root development and root-microbe interactions in woody plants. Here, we developed the RMI-chip, a simple microfluidic setup in which *Populus tremuloides* (aspen tree) seedlings can grow for over a month, allowing continuous microscopic observation of interactions between live roots and rhizobacteria. We find that the colonization of growing aspen roots by *Pseudomonas fluorescens* in the RMI-chip involves dynamic biofilm formation and dispersal, in keeping with previous observations in a different experimental set-up. Also, we find that whole-cell biosensors based on the rhizobacterium *Bacillus subtilis* can be used to monitor compositional changes in the rhizosphere but that the application of these biosensors is limited by their efficiency at colonizing aspen roots and persisting. These results indicate that functional imaging of dynamic root-bacteria interactions in the RMI-chip requires careful matching between the host plant and the bacterial root colonizer.

## Introduction

The plant microbiome plays an important role in the rhizosphere (1–3). Virtually all plant tissues host microbes that can act as symbionts, commensals, or pathogens. Interactions between plant and microbes can be beneficial, neutral, or harmful and directly influence plant growth and productivity (2,4,5). Plant-growth-promoting (PGP) rhizobacteria are bacteria that exert beneficial effects on plants through direct or indirect interactions with the roots (6,7). PGP bacteria have the potential to increase the availability of soil nutrients to the plant, produce metabolites such as plant hormones, elicit plant systemic defenses, and increase plant resistance to biotic and abiotic stresses (8,9). In return, the plant provides photosynthetically-derived carbon, such as sugars and organic acids that are consumed by rhizosphere microorganisms as well as a wide range of molecular compounds acting as environmental signals for the root microbiota. Microbes attach to the root surface and form micro-colonies that can eventually grow into larger biofilms. The formation of biofilms at root surfaces was proposed to be part of the cellular PGP activities of beneficial rhizobacteria (10).

Understanding the complex interactions between plant roots and microbes requires the ability to track their dynamics at high spatial and temporal resolution. Real time monitoring of dynamic root-microbe interactions at cellular resolution is now possible using microfluidics approaches coupled with advanced live imaging microscopy. Microfluidic platforms provide a powerful approach to evaluate the responses of growing plant cells to external perturbations (e.g., nutrients, media flow, temperature, hydrodynamics, light, and stressors) at throughputs higher than with conventional methods using pots or plates, and in precisely controlled environments. Multiple microfluidics devices such as “Plant on a chip” (11), RootChip (12), RootArray (13), TipChip (14), and PlantChip (15) were developed to study various aspects of the cell biology of *Arabidopsis thaliana*, including gene expression, cell biomechanics, cellular mechanism of growth and cell division (reviewed in (16)). The PlantChip device enables the continuous monitoring of phenotypic changes at the cellular level and also at the whole plant level, including seed germination and root and shoot growth (hypocotyls, cotyledons, and leaves) (15).

Fewer studies have used microfluidic devices to visualize the interactions of *Arabidopsis* roots with pathogenic or beneficial microorganisms. Using the plant-in-chip platform, visualization of the attack of *Arabidopsis* roots by pathogenic nematodes and oomycetes motile spores revealed some physiological changes taking place in the host plant and the pathogen during the attack (17). Recently, a microfluidic device to track root interactions system (TRIS) revealed a distinct chemotactic behavior of the bacterium *Bacillus subtilis* toward the root elongation zone and its rapid colonization, and allowed real-time monitoring of bacterial preference between roots from various *Arabidopsis* genotypes (18). Recent study investigated the spatiotemporal dynamics of colonization of *Arabidopsis* roots by PGP bacterial species from the P. deltoids rhizosphere over 4 days {Retterer, 2018 #59}. To date, studies of plant roots and root-microbe interactions (RMIs) using microfluidics have been focused on *A. thaliana*, an annual herbaceous plant model that can complete is entire life cycle in 6 weeks, and grows a single primary root that later produces smaller lateral roots. However, there is a need to study root development and root-microbe interactions for other plants, including woody perennial plants such as trees.

Here, we describe a microfluidic device, called RMI-chip, that enables the direct visualization of root-microbe interactions taking place at early stages of tree seedling growth. We studied the interactions of *Populus tremuloides* (trembling as pen tree) with the bacterium *Pseudomonas fluorescens*. These interactions are biologically relevant in nature, as *P. fluorescens* is abundant in the rhizosphere of *Populus* trees (1,19–21), and exhibits functionality in laboratory assays. We have shown that *P. fluorescens* promotes the growth of aspen seedlings (22), colonizes aspen seedling roots by forming dense and dynamic biofilms (23), and modulates expression of anti-fungal defense response genes in roots of aspen seedlings (24). The RMI-chip device was designed to accommodate aspen seedling growth for periods up to 5 weeks, and to enable direct observation of root growth and its dynamic colonization by *P. fluorescens* biofilms with high spatiotemporal resolution. The RMI-chip was also used to monitor the growth of rice seedling roots and detect the production of reactive oxygen species (ROS) by the root using engineered *Bacillus subtilis* strains as biosensors. We find that in the RMI-chip interactions between host plants and bacterial species are specific, consistent with ecological observations and with colonization profiles observed in other experimental systems, and that formation of bacterial biofilms on root surfaces is needed for persistent colonization.

## Materials and Methods

### Seeds and growth media

Seeds of Populus tremuloides Michx. were obtained from the National Tree Seed Center, Natural Resources Canada, Fredericton NB, Canada. The rice seeds were obtained from Baker Creek Heirloom Seed Co. (Mansfield, MO, USA). Phytoblend was purchased from Caisson Laboratories, Inc. (Smithfield, UT, USA). All other chemicals were purchased from Sigma-Aldrich (St. Louis, MO, USA).

### RMI-chip and humidity chamber design

The chips were designed with AutoCAD software (Autodesk) and were fabricated via soft lithography at the Scientific Device Laboratory (Des Plaines, IL, USA). Briefly, SU-8 2025 photoresist (MicroChem, Westborough, MA, USA) was used to make molds on a 4-inch silicon wafer. The two-part silicone elastomer (SYLGARD 184, Thermo Fisher Scientific, Waltham, MA), the silane precursor and curing agent were mixed 10:1, degassed, poured on silicon wafer mold, degassed and baked at 65 °C for 4 hours. The PDMS elastomer was punched and bonded to a 48 x 65mm, thickness No. 1 cover glass (Ted Pella Inc., Redding, CA, USA). RMI-chip devices were used without Aquapel treatment. The humidity chambers and microscope stage adapters were fabricated from PMMA thermoplastic via laser cutting (Ponoko, Oakland, CA, USA) and assembled with acrylic adhesive (Weld-On, Compton, CA, USA). Tubing management and other holder accessories were 3D printed using PLA. The AutoCAD files (ESI) of the RMI-chip design, the SVG files used for laser cutting, and the SCAD and STL files used for 3D printing are available in the online supporting material.

### Fluorescent-labelled and biosensor bacterial strains

The mNeonGreen (mNG)- and dsRed-labelled *P. fluorescens* SBW25 strains harbored an environmentally stable plasmid derivative that constitutively expresses the dsRed or mNG fluorescent proteins, as described (25). B. subtilis strains labelled with the mCherry fluorescent protein were constructed by transferring the genetic construct P_pen_-mCherry::kan from *B. subtilis* MMB1023 (26) into the NCIB3610 background by SPP1-mediated phage transduction. The resulting NCIB3610-mcherry strain, which expressed constitutively mCherry, was used as a recipient for the transfer of sensory genetic elements. A xylose-responsive genetic module was constructed by inserting a P_xylA_ BioBrick DNA block (PCR-amplified from plasmid pBS1C3-PxylA) into plasmid pBS1C (27) to form pBS1C-PxylA, which was then used as recipient for the insertion of the *gfp_sp_* gene from pRD111 (28), using Spe1 and Pst1 restriction sites. In the final plasmid pBS1C-P_xylA_-gfp_sp_, the *gfp_sp_* gene is under control of the xylose-inducible P_xylA_ promoter, linked with a chloramphenicol resistance gene (cat) and flanked by the right and left arms of the *B. subtilis amyE* gene. The P_xylA_-GFP::cat module was integrated in the *amyE* gene of *B. subtilis* strain 1A976 (SKC6) by transformation of the competent cells. The module was then finally transferred in the NCIB3610-mCherry strain by SPP1-mediated phage transduction. A ROS-responsive *B. subtilis* strain was constructed by SPP1 transduction of the genetic module P_katA_-*gfp_sp_::spc* from strain 1A1010 (Bacillus Genetic Stock Center, http://www.bgsc.org), where P_katA_ is the promoter controlling the expression the catalase gene, in the NCIB3610 strain for integration at *amyE* locus. Transductants were selected for resistance to spectinomycin.

### Cultivation of aspen and rice seedlings in the RMI-chip

*Populus tremuloides* Michx. seeds were cold stratified in Milli-Q water at 4°C for 2 to 14 days. Seeds were then surface sterilized by prewashing briefly in 1% Tween-20 and incubated in pH-reduced (10 mM HCl) 0.1M sodium hypochlorite. After 4 minutes of incubation the seeds were rinsed 8-10 times with sterile Milli-Q water and incubated in sterile Milli-Q water overnight in the dark at room temperature.

The following day the seeds were spread onto 1% agar Johnson’s plate (4 mM KNO_3_, 2 mM Ca(NO_3_)_2_, 4 mM NH_4_NO_3_, 0.5 mM KH_2_PO_4_, 1 mM MgSO_4_, 440 µM KCl, 250 µM H_3_BO_3_, 20 µM MnSO_4_, 20 µM ZnSO_4_, 5 µM CuSO_4_, 5 µM NaMoO_4_, 5 µM CoCl_2_, 200 µM Fe,Na-EDTA, pH 5.6, 1% Phytoblend). The seeds were incubated under low to medium light on a 16hr light/8hr dark period, maintaining the temperature below 30°C. Once germinated, seedlings with hypocotyls shorter than 5 mm were transferred into cut 200 µL pipette tips filled with 1% Johnson’s agar to let the root elongate gravitropically. For a typical experiment, 20-25 pipette tips were inserted into Johnson’s agar (1%) plates at a ~45° angle and incubated under the same conditions (Fig. S1). Root growth was checked periodically. Six seedling-containing tips with roots grown close to the end of the tips were mounted into the RMI-chip, which was incubated at a ~45° angle to facilitate root infiltration.

De-hulled rice seeds (*Oryza sativa* ‘Diamante-INIA’) were prewashed briefly in 1% Tween-20, followed by incubation in pH-reduced (40 mM HCl) 0.4M sodium hypochlorite solution on a rocker for 15-20 minutes. The seeds were rinsed 7 times with Milli-Q water, plated on 1% Johnson’s agar, and incubated at 30°C in the dark for 48-72 hours. When germinated and the radical protrusions reached 3-6mm, the seedlings were planted directly into cut 200 µL pipette tips filled with Johnson’s solution already inserted into the RMI-chip.

Following hypochlorite sterilization, all seed and seedling manipulations were performed in a laminar flow hood, including mounting in the RMI-chip. Syringes, tubing, and adaptors were autoclaved prior to connection to the chip to maintain sterility during the extended continuous perfusion.

### Bacterial inoculation of the RMI-chip

Bacterial species were cultivated overnight in LB at their respective optimal temperature (i.e. 28°C for *P. fluorescens*, 37°C for *B. subtilis*) in the presence of the appropriate antibiotic selection. Overnight cultures of *P. fluorescens* SBW25-mNG and-DsRed strains were centrifuged, rinsed in PBS, and re-suspended in PBS to OD_600_=4. A 10μl aliquot of cell suspension was used for inoculation of aspen roots. Overnight cultures of *B. subtilis* biosensor strains were diluted 100-fold and grown in LB up to mid-exponential phase (OD_600_=0.4), rinsed with PBS and likewise adjusted to OD_600_=4 for the inoculation of the roots of aspen and rice seedlings. After overnight incubation without flow, excess bacteria were washed away with Johnson’s solution at a flow of 0.2 µL/min for 2 hours, followed by continuous perfusion at 0.02 µl/min. The outlet ports were connected to 200 µL pipette tips during inoculation, excess liquid was removed, and ports were connected to independent lines to prevent cross-communication between the 100 µm x 800 µm channels (Fig. 1a) and with the external Johnson’s solution overlay during perfusion. Note that in theory, the RMI-chip can be washed and autoclaved after first use making the second set of 6 independent channels available for a new experiment. In this study, RMI-chips were used only once.

**Fig. 1.**
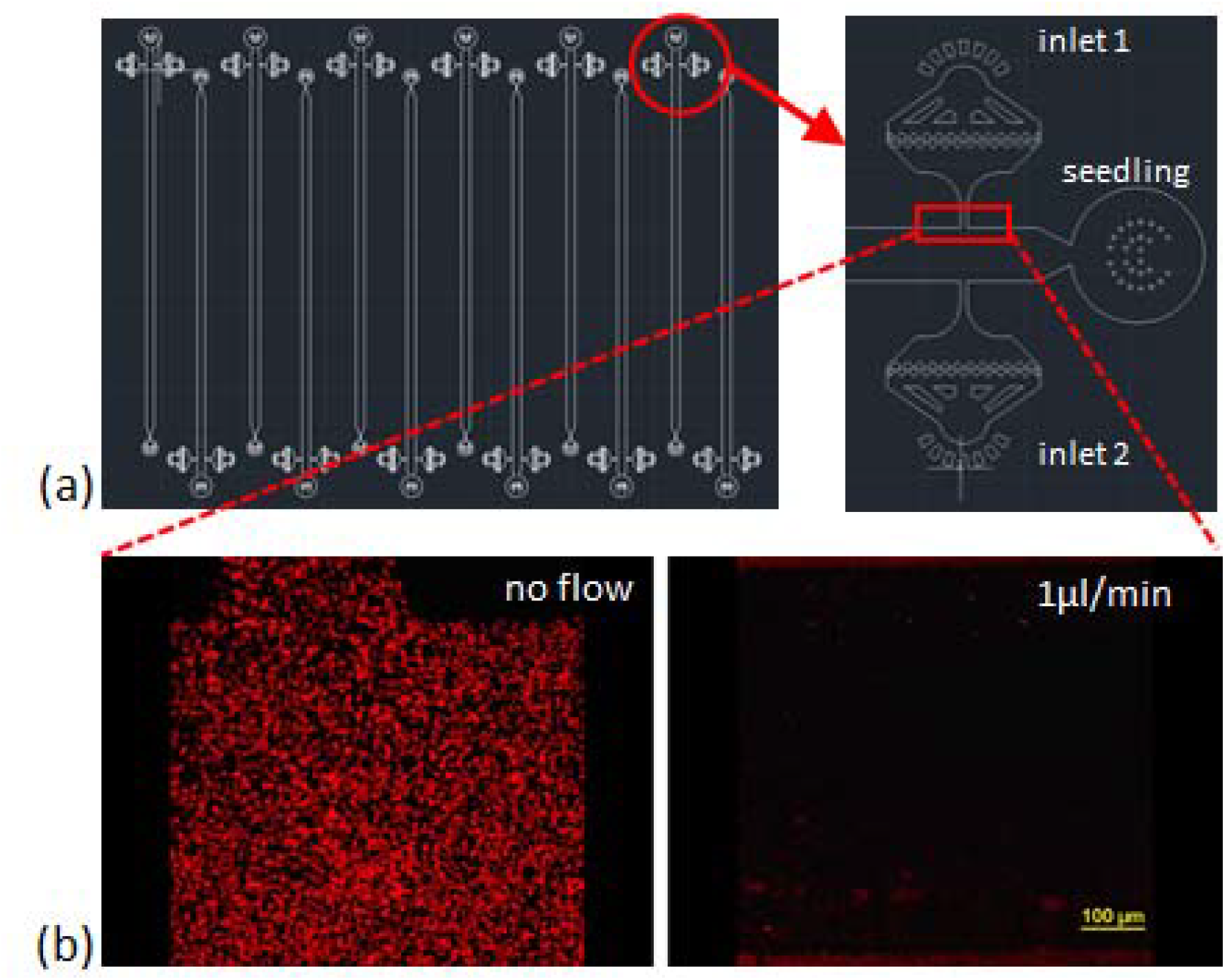
RMI-chip design. (a) The CAD design of the RMI-chip. A single growth channel is 36 mm long, 100 µm high, and 800 µm wide. The media and inoculation inlets are equipped with filters to avoid occlusions during week-long experiments. Design files are available in the online supporting materials. (b) Flow dynamics. Fluorescently labelled latex beads were used to measure media flow in the empty RMI-chip. The root entrance and the inoculation inlets were plugged with stopper pins to mimic operational flow regime of the chip. The latex beads suspended in Johnson were injected and the chip images, followed by perfusion with Johnson solution at 1µL/min. The latex beads were washed from the upstream chambers and the velocity of the beads were averaged. The measured and calculated values agreed. Laminar flow was observed under the microscope.

### Imaging experiments

An inverted Nikon C2+ laser-scanning confocal microscope was used for imaging experiments (Nikon, Melville, NY, USA). An Eclipse Ti-E inverted microscope equipped with perfect focus system, an automated stage, and with 10x, 20x, and 100x objective lenses (CFI Plan Fluor 10X, NA 0.3, WD 16 mm; CFI Plan Apochromat Lambda 20x, NA 0.75, WD 1.00 mm; CFI Plan Apo Lambda 100x, NA 1.5, WD 130 um, respectively) was used for single image, time series, and z-stack acquisitions. Laser illumination emission at 488 nm coupled with a 525/50-nm excitation filter was used to capture mNeonGreen fluorescence, and laser illumination at 561 nm coupled with a 595/50-nm excitation filter was used to capture dsRed (*P. fluorescens*) or mCherry (*B. subtilis*) fluorescence. The transmitted light was also detected to image bacterial colonization in the context of the root structure. A custom holder was designed to provide access of the objective to the RMI-chip and imaging chamber.

## Results and discussion

### RMI-chip design optimized for aspen seedling growth

The RMI-chip was designed to observe the growth and rhizobacterial colonization of aspen seedling roots over extended periods of time. The aspen root system consists of a taproot from which smaller branch roots emerge. When a seed germinates, the first root to emerge is the primary root which develops into the taproot. With seedlings grown into agar-filled pipette tips (Fig. S1a), we observed that the primary root quickly branched after exiting the tip (Fig. S1b, Fig. S2a). Thus, seedlings were inserted into the RMI-chip shortly before the primary root reached the end of the pipette tip. In early studies, we observed that although the primary root readily entered the RMI-chip circular chamber (Fig. 1a), a substantial fraction of the roots did not continue into the linear growth channel, instead growing in circles and causing the seedling to wither quickly. This problem was resolved by growing the seedlings semi-gravitropically in a chip tilted at a 45° angle, submerged in Johnson’s solution for up to a week, until the root tip reached the media inlets in the growth channel (Fig. S2c). Under these conditions, most primary roots entered the growth channel, at which point the RMI-chip was placed horizontally in a humidity chamber and continuous media flow was supplied (Fig. 2). Regular exchange of the nutrient solution surrounding the RMI-chip and the maintenance of water-saturated paper used to provide the required humidity in the growth chamber, were crucial to obtain healthy seedling growth.

**Fig. 2.**
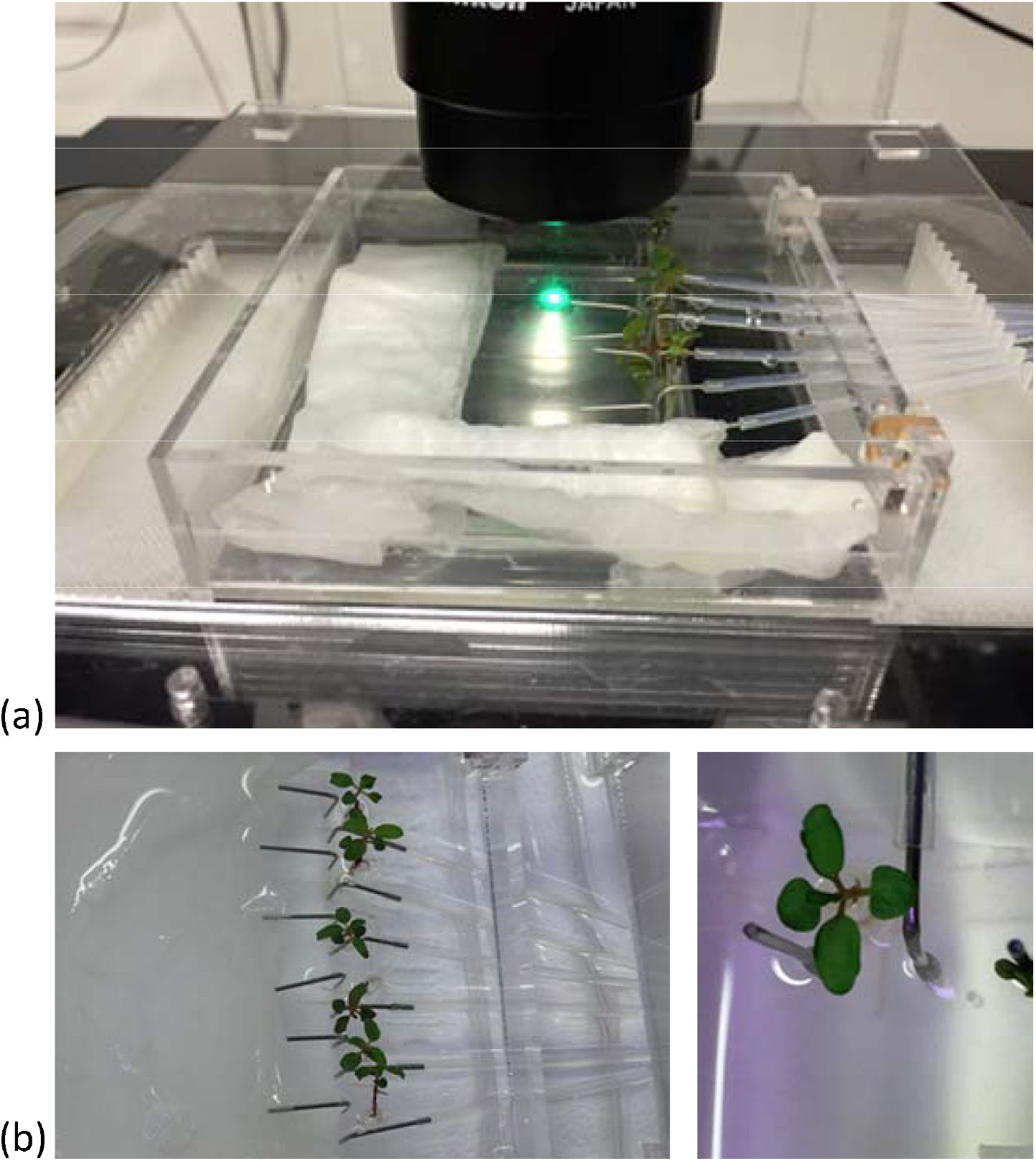
Imaging setup. (a) The imaging setup with humidity chamber and 3D printed stage adaptor. The RMI-chip has an inlet for media, a separate port for inoculation, and a wide channel to accommodate the tree seedling root without interfering with constant media flow. (b) The continuous flow incubation chamber enables the cultivation of six seedlings in parallel (left). Details of an aspen seedling (right).

The RMI-chip has 6 independent channels with one inlet for the seedling root, two dedicated inlets with filters for media and bacterial inoculation, and one common outlet (Fig. 1a). The features are generated via soft lithography on a silicon wafer, which is then used as a mold to produce polydimethylsiloxane (PDMS) slabs. The PDMS is bonded to a large thin microscope coverslip providing a means for imaging at cellular resolution using a confocal microscope. Different channel depths were tested for aspen seedling growth. Although aspen primary root can grow into an 80 µm x 80 µm channel in a submerged microfluidic device (Fig. S2b), the root occupied the entire channel precluding flow experiments. Therefore, we selected a channel width of 800 µm and fabricated channels with depths of 100 µm and 400 µm. While aspen seedlings grew similarly in both devices, the 400 µm deep channel did not position the root close to the glass coverslip, making it suboptimal for microscopic observations using objectives with standard working distances (Fig. 2). Our final design has growth channels 100 µm deep, 800 µm wide and 36 mm long, which can easily accommodate aspen primary roots and enable root growth under constant perfusion. Flow characteristics of the 400 µm x 800 channel were measured with fluorescently labelled styrene beads (Fig. 1b). Without perfusion, slight bulk material movement was detected. At 1 µL/min, laminar flow was observed with a measured 50 µm/s bead velocity (Supplementary video 1).

It can be challenging for plant roots to grow under constant flow. For example, without flow in a growth channel, root border cells and mucilage can be observed at the root tip (Fig. S2c, Fig. S3). These cells and mucilage can be washed away at high flow rates (e.g., 2 µL/min). Nevertheless, nutrient flow is needed to maintain seedling growth in the RMI-chip. Therefore, we determined a minimal flow rate of nutrient solution that did not affect root growth, preserving root morphology, including root cap, root hairs and mucilage, while keeping out air bubbles. After several iterations, each including at least 4 seedlings, the minimal flow rate was found to be 0.02 µL/min. As each channel is connected to an individual 1 ml syringe, this flow rate provides enough media for up to 5 weeks. Under these conditions, the calculated average laminar flow is 4 µm/s and the media is replaced every ~14 minutes in each channel. We found that aspen seedlings exhibited a high heterogeneity in primary root growth with an average growth rate of 1 ± 0.9 mm/d (n=10). This heterogeneity may be due to the physical stress imposed on the seedling by the RMI-chip confined space combined with our use of open pollinated seeds which have different genetic makeups. Generally, only 4 or 5 seedlings out of 6 mounted in the RMI-chip typically sustained growth over a 5-week period. On average, aspen primary roots grew much slower than *Arabidopsis* primary roots reported to grow at 3.7 mm/d in the RootChip (12).

### Continuous imaging of the RMI-chip

Confocal Laser Scanning Microscopy was used to perform spatial and temporal studies of the aspen root growth. Because of the repeated microscopy observations, a flexible system was required which was compatible with the transfer of the chip between the growth chamber and microscope stage. A modular chamber was designed to overcome this technical challenge and to prevent water evaporation during cultivation and imaging. An innermost imaging chamber was built to provide humidity during the repeated 1-hour imaging experiments, and a medium-size chamber allowed submersion of the RMI-chip in media and its transport between incubation chamber and microscope. A six-channel syringe pump was connected with Teflon™ FEP tubing to the chip and transferred with the chip during imaging experiments in order to provide continuous flow of nutrients, even during long imaging experiments (Fig. S4). Finally, a large closed chamber that can be filled with water was built to provide humidity during long-term growth. In addition, a holder compatible with the imaging chamber was fabricated for the Nikon Ti-2 microscope (Fig. S4). The humidity chambers and microscope stage adapters were fabricated from PMMA thermoplastic. 3D printed inserts were fabricated to manage the tubing during imaging experiments (design files are available in the online supporting material). With this system, the RMI-chip was continuously perfused with Johnson’s solution flowing at 0.02 µL/min, including during the lengthy observations under the microscope, thus minimizing the disturbance of seedling growth.

### Growing rice seedlings in the RMI-chip

We tested the compatibility of the RMI-chip with other plant species such as rice. Rice seedling roots could be introduced easily into the chamber without an agar tip by directly transplanting the seedlings within days after germination. Healthy root growth was observed under continuous perfusion with Johnson’s solution at a 0.02 µL/min flow rate (Fig. S5). Under these conditions, we observed fast root growth where the root tip reached the end of growth chamber after 4 days on average. As for aspen, variability between seedlings was observed, some root tips reaching RMI-chip end after 3.5 to 5 days. Thus, rice root growth rate is approximately 10 times faster than aspen root growth under the same conditions. Unfortunately, the fast growth rate of rice seedlings limits observation time to 3-5 days. Once the root tip reaches the outlet, it blocks the chamber and cuts off nutrient flow.

### Colonization of aspen seedling root by *P. fluorescens*

Aspen seedlings grown for 8 days in the RMI-chip were inoculated with mNeonGreen-labelled *P. fluorescens* SBW25 by injection of approximately 2 × 10^9^ cells through an inlet port (Fig. S6a). After incubation for 16 hours without flow, the flow was restored at 0.2 µL/min for 2 hours to wash away excess bacterial cells, then set at 0.02 µl/min for continuous perfusion. Under these conditions, a small number of SBW25 cells remained associated with the lower part of the root (Fig. S6b). Importantly, the only source of carbon available for bacterial growth in the RMI-chip are the sugars and organic acids photosynthetically produced by the plant. Five days after inoculation, actively dividing SBW25 cells were colonizing the intercellular spaces between root epidermal cells of the root cortex, and, to a lesser extent, the root surface (Fig. S7). Although this preferential colonization of crevices on the root surface may be caused by the flow in the RMI-chip, it is consistent with our previous observations in a vertical plate system, where SBW25 first colonizes the cell interstitial spaces along the aspen root cortex before forming a variety of bacterial assemblies that range from microcolonies to highly structured biofilms (23).

Next, we observed aspen primary root colonization by a 1:1 mixture of two *P. fluorescens* SBW25 derivatives labelled with distinct fluorescent proteins, mNeonGreen and dsRed. Three days after inoculation, we observed discrete patches of red and green cells with almost no area of mixed colors (Fig. 3a). This finding indicates that bacterial cells divide and form expanding patches in the intercellular spaces and on the root surface, ruling out that those patches are formed by random deposition and aggregation of cells within the root crevices. Thirteen days after inoculation, bacterial patches were longer and formed dense cell assemblies on the primary root surface (Fig. 3b). Close-up examination revealed mature biofilms resulting from clonal growth of green and red bacterial populations. These biofilms exhibited channel-like void spaces, which are a hallmark of mature biofilms (29). Although cell density and presence of small aggregates in the pure cultures prior to co-inoculation could play a role in determining the colonization patterns (30), the quasi-absence of intermixed biofilms is indicative of a competition between the green and red strains. These strains have an identical genome but differ by the expressed fluorescent protein, possibly creating differences in cell fitness and some competition. Notably, we observed that a variety of SBW25 cell assemblies coexisted with biofilms on the root surface (Fig. 4a). These assemblies included clusters of somewhat regularly spaced individual cells, clusters of cell doublets which are presumably dividing, microcolonies and mature biofilms (Fig. S8). These assemblies may reflect the different stages of biofilm formation, where cells attach to the root surface, divide, and over time form more compact assemblies maturing into biofilms. Intriguingly, cells packed within a biofilm appeared to have a round coccoid shape in contrast with the normal rod-like morphology of individual SBW25 cells (Fig. 3a, Fig. 4a). Confocal 3D image reconstruction and side projection of the 3D volume (Fig. 4b) revealed that the coccoid shape was only apparent and resulted from the top view of vertically arranged bacterial cells within the biofilm. This finding corroborates our previous observation of vertical arrangements of SBW25 cells within biofilms and mucilage formed on the roots of aspen seedlings growing in a vertical agar plate (23).

**Fig. 3.**
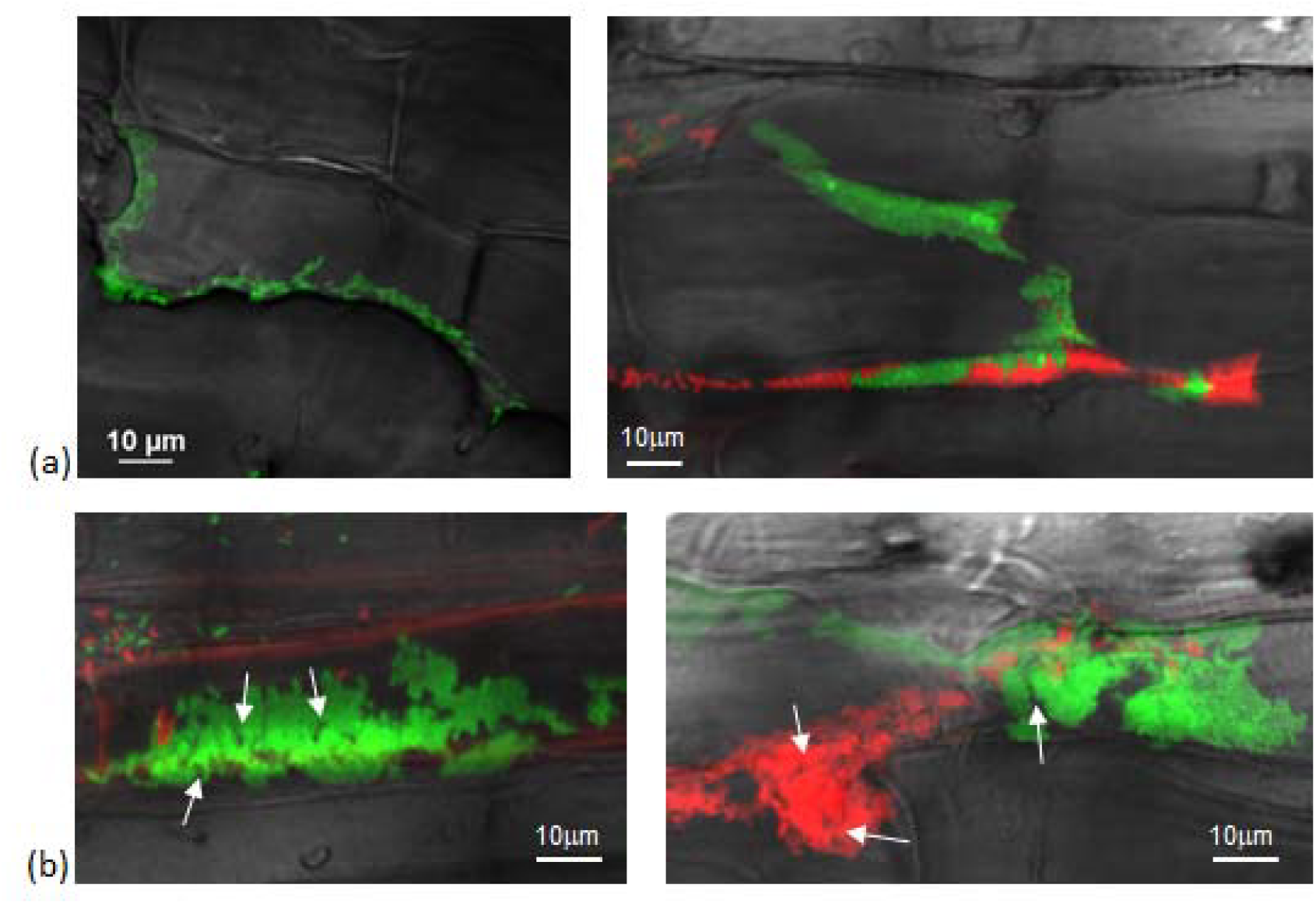
Spatial and temporal colonization of aspen primary roots by *P. fluorescens* in the RMI-chip. *P. fluorescens* SBW25 strains expressing dsRed or mNeonGeen were co-inoculated on aspen roots growing in the RMI-chip. (a) After 3 days, bacterial cells colonize aspen root surface by filling intercellular spaces (left panel) and adhering to plant cell surfaces (right panel). (b) After 13 days, SBW25 cells formed spatially segregated red and green long patches of cells that correspond to dense biofilm-like assemblies. Typical of mature biofilms, some internal void spaces and channels are indicated by arrows.

**Fig. 4:**
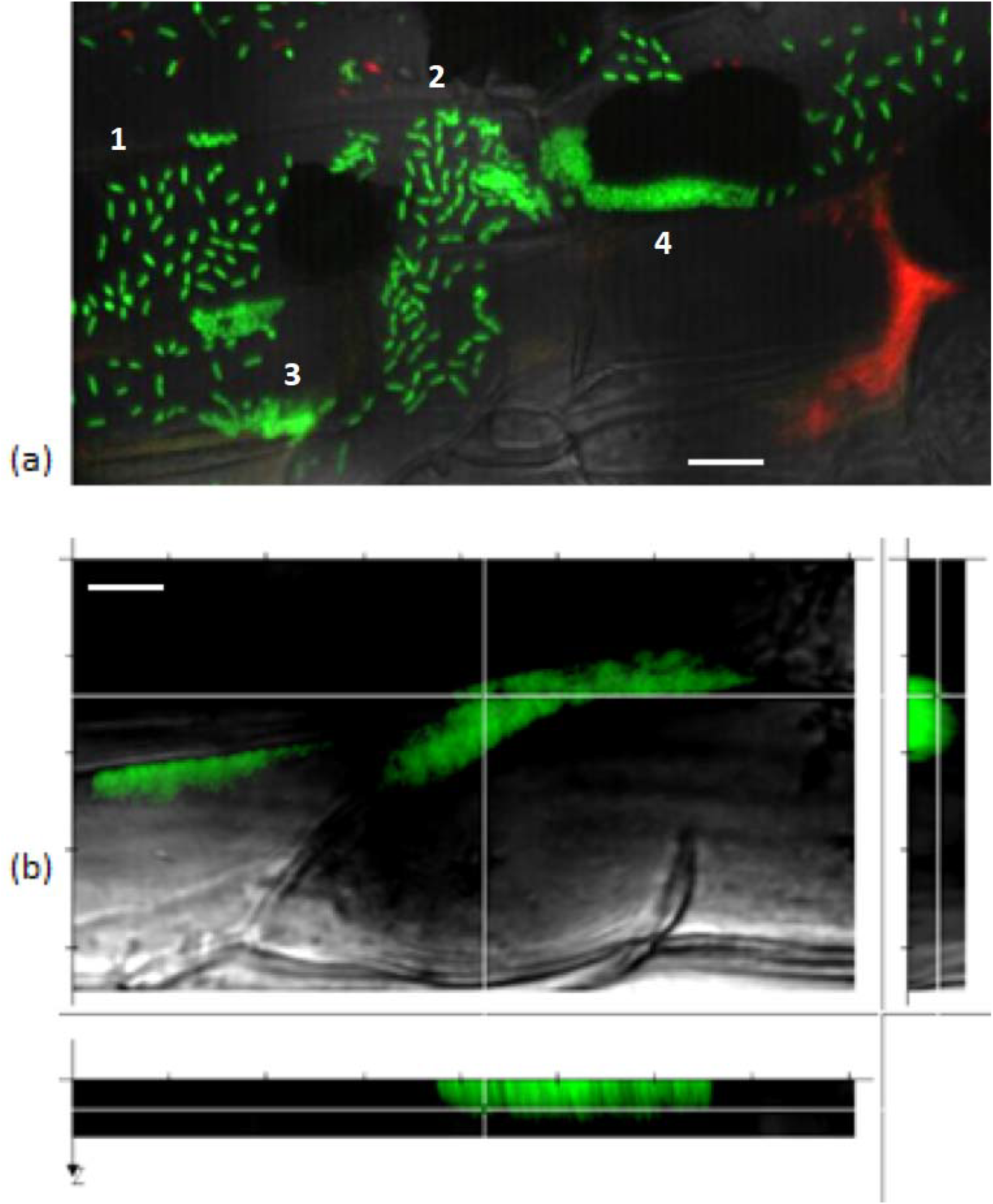
A gallery of bacterial assemblies on the root surface. (a) Image showing diverse assemblies of green SBW25 cells on the root surface: 1, individual cells; 2, cell doublets; 3, microcolonies; and 4, biofilms. (b) Projection of a reconstructed 3D volume with side views reveals a vertical alignment of cells within the biofilm. Scale bar indicates 10 µm length.

After 30 days in the RMI-chip, fluorescent SBW25 cells became rare on the root surface and were preferentially found as micro-colonies located within the spaces between root epidermal cells (Fig. S9). Overall, the dynamic of root colonization by SBW25, which includes active growth and colonization of the root during the first two weeks, formation of mature biofilms associated with a variety of other cell assemblies, and dispersal of these biofilms and assemblies after 5 weeks, appears to be remarkably comparable in the RMI-chip and vertical plate assay, the latter being a very different set-up where seedling roots grow on a nutrient agar surface (23). This similarity suggests that the observed dynamics of cell assemblies are not primarily shaped by the flow in the RMI--chip but rather reflect an intrinsic colonization behavior of aspen primary root by *P. fluorescens* SBW25. In addition, novel cell assemblies such as clusters of somewhat regularly spaced individual cells and cell doublets were observed uniquely in the RMI-chip.

### RMI-chip and rhizobacterial biosensors to monitor dynamic root exudation

In the RMI-chip, the carbon sources available for bacterial growth are provided by root exudates. These exudates are compositionally complex and contain ions (i.e. H^+^), inorganic acids, oxygen, water, and multiple carbon-based compounds which include amino acids, organic acids, sugars, phenolics and an array of secondary metabolites, and high-molecular weight compounds like mucilage and proteins (31). Root exudates are key mediators of interactions with microbes in the rhizosphere and root exudation is a regulated process which responds to biotic and abiotic stresses to the plant (31). We assessed whether whole-cell biosensors, which are engineered rhizobacteria that can express a fluorescent protein in response to the presence of a specific metabolite or environmental stressor, could be used in the RMI-chip to monitor dynamic changes in the composition of plant exudates.

We used *Bacillus subtilis*, which colonizes *Arabidospis* roots (18) and is used as biocontrol agent to colonize and protect various herbaceous plants (32), to develop biosensor strains. We selected two well characterized promoters, one responding to the presence of xylose (33), a sugar abundant in root exudates, and the other one responding to the presence of Reactive Oxygen Species (ROS) (34), which are normal products of plant cellular metabolism and act as second messengers in plant responses to various environmental stresses (35). The biosensors expressed the green fluorescent protein (GFP) conditionally to the presence of xylose or ROS and also expressed constitutively the red fluorescent protein mCherry, which serves to label the cells (see Experimental methods).

Aspen primary roots were inoculated with *B. subtilis* biosensors in the RMI-chip, incubated overnight without flow, and upon restoration of flow, the root was observed using confocal laser microscopy. We found that, as with *P. fluorescens*, most cells were rapidly washed away from the root surface. However, biosensor cells were very sparse on the root after 3 days and could not be observed after 5 days. The use of a low flow (0.02 µL/min) throughout inoculation of the RMI-chip did not enhance the number of biosensor cells remaining attached to the primary root, reflecting an inability of *B. subtilis* to colonize aspen primary root under these conditions. This is in contrast with our observations with *P. fluorescens* and may reflect the fact that *B. subtilis* is not a natural colonizer of aspen roots. To circumvent this problem, observations were done in a stopped-flow experiment limited to 5 days, after which the aspen seedling starts to wither. Even in the absence of flow, Bacillus cells did not colonize but remained loosely attached to the aspen root surface. Whereas biosensor cells did not express GFP after inoculation (Fig. 5 a), strong GFP signals were detected in biosensor cells remaining attached to the aspen root after 5 days of incubation, indicating the production of xylose (Fig. 5b) or ROS (Fig. 5c) by the root. Exposure of the ROS biosensor to a rice primary root under the same RMI-chip conditions (i.e., no flow) resulted in a robust colonization of the root and in high levels of GFP after 3 days (Fig. 5d). This finding indicates that high levels of ROS are produced by rice primary root under these conditions. Together, these results provide a proof-of-concept that bacterial biosensors can be used in the RMI-chip as a way to investigate the dynamic chemical crosstalk between root and rhizobacteria.

**Fig. 5.**
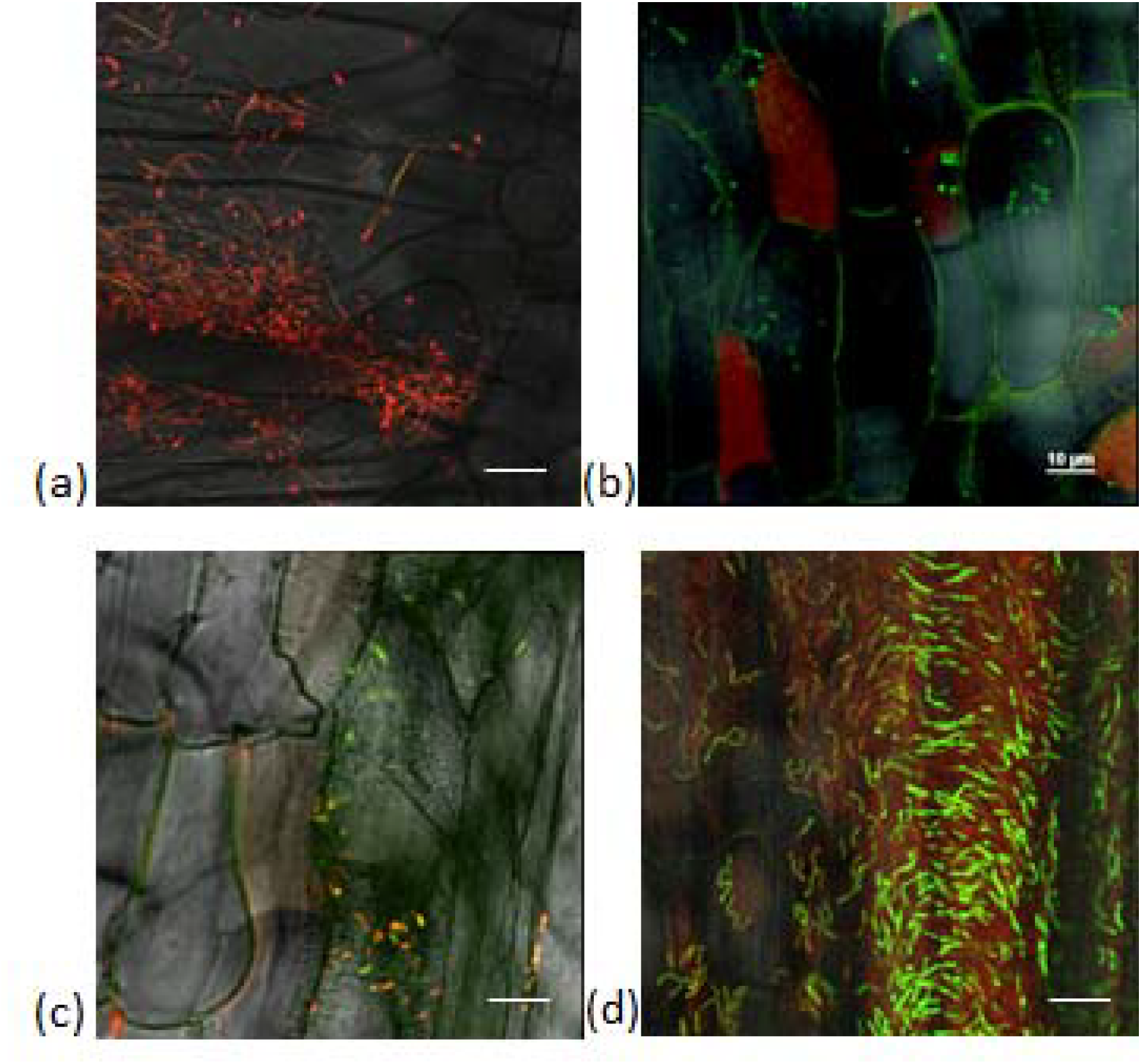
*B. subtilis* biosensors exposed to aspen root (panels a-c) and rice root (panel d) in the RMI-chip. (a) Among the root hairs, the xylose biosensor cells appear red from constitutive mCherry expression with a very faint GFP (green) signal detected in some cells. Similar results were obtained with the ROS biosensor (not shown). (b) Five days after inoculation, rare attached biosensor cells overwhelmingly express GFP, indicating the presence of xylose in root exudates. Note the plant tissue autofluorescence causing cell outlines to be green and some plant cells to be entirely red. (c) After 5 days, most of the rare ROS biosensor cells expressed GFP, indicating the presence of ROS produced by the root. Plant tissue autofluorescence is also detected. (d) Three days after inoculation of a rice seedling, the *B. subtilis* ROS biosensor displayed robust colonization of the root with a strong expression of GFP, indicating production of ROS by the rice root. Scale bar indicates 10 µm length.

We also inoculated rice seedlings with *P. fluorescens* SBW25 to test for ability to colonize in the RMI-chip. Repeated trials showed that SBW25 cells were rapidly washed away from the root surface and remained barely detectable under slow nutrient flow after one day (data not shown), suggesting an inability of *P. fluorescens* SBW25 to colonize rice primary root under these conditions. This result was unexpected as rice was shown to host *Pseudomonas* endophytes (36). These observations clearly indicate that biosensors need to be developed from bacterial isolates that efficiently colonize the roots of the plant host of interest. What may seem to be a caveat for the development of biosensors is actually grounded in ecology and evolution, as root exudates from a particular plant species are known to maintain and support a highly specific diversity of microbes in the rhizosphere of this plant (31).

## Conclusions

In summary, we report the design and fabrication of a microfluidic device and accessories that enable the cultivation of aspen seedlings under constant nutrient flow and the study of root-microbe interactions through repeated functional imaging during a 5 week long experiment. The device enabled imaging at single bacterial cell resolution during the various phases of aspen colonization by *P. fluorescens* SBW25. Different biofilm morphologies and diverse bacterial assemblies were observed over time, emphasizing the need for high-resolution imaging to understand colonization patterns and strategies and their relation to the local root environment. *B. subtilis* whole-cell biosensors provided a means for functional imaging of bacterial cells actively consuming xylose from root exudates and actively responding to ROS, an environmental stressor produced by the root. However, the use of these biosensors to image compositional changes in the root environment was limited by the inability of *B. subtilis* biosensor to colonize and persist on aspen roots, suggesting that whole-cell biosensors should be built from naturally colonizing bacteria. The current design of the RMI-chip can be used for the long-term observation of slow-growing plants, or can be modified to study faster growing plants as well.

## Supporting information

Flow dynamics in RMI-chip

Aspen root tip in RMI-chip

oligos and strains used

ESI

## Conflicts of interest

There are no conflicts to declare.

## Acknowledgements

This work was supported by funding through the Biological Systems Science Division, Office of Biological and Environmental Research, Office of Science, U.S. Dept. of Energy, under Contract DE-AC02-06CH11357.

