## Supplementary material for "Functional imaging of microbial interactions with tree roots using a microfluidics setup": ESI

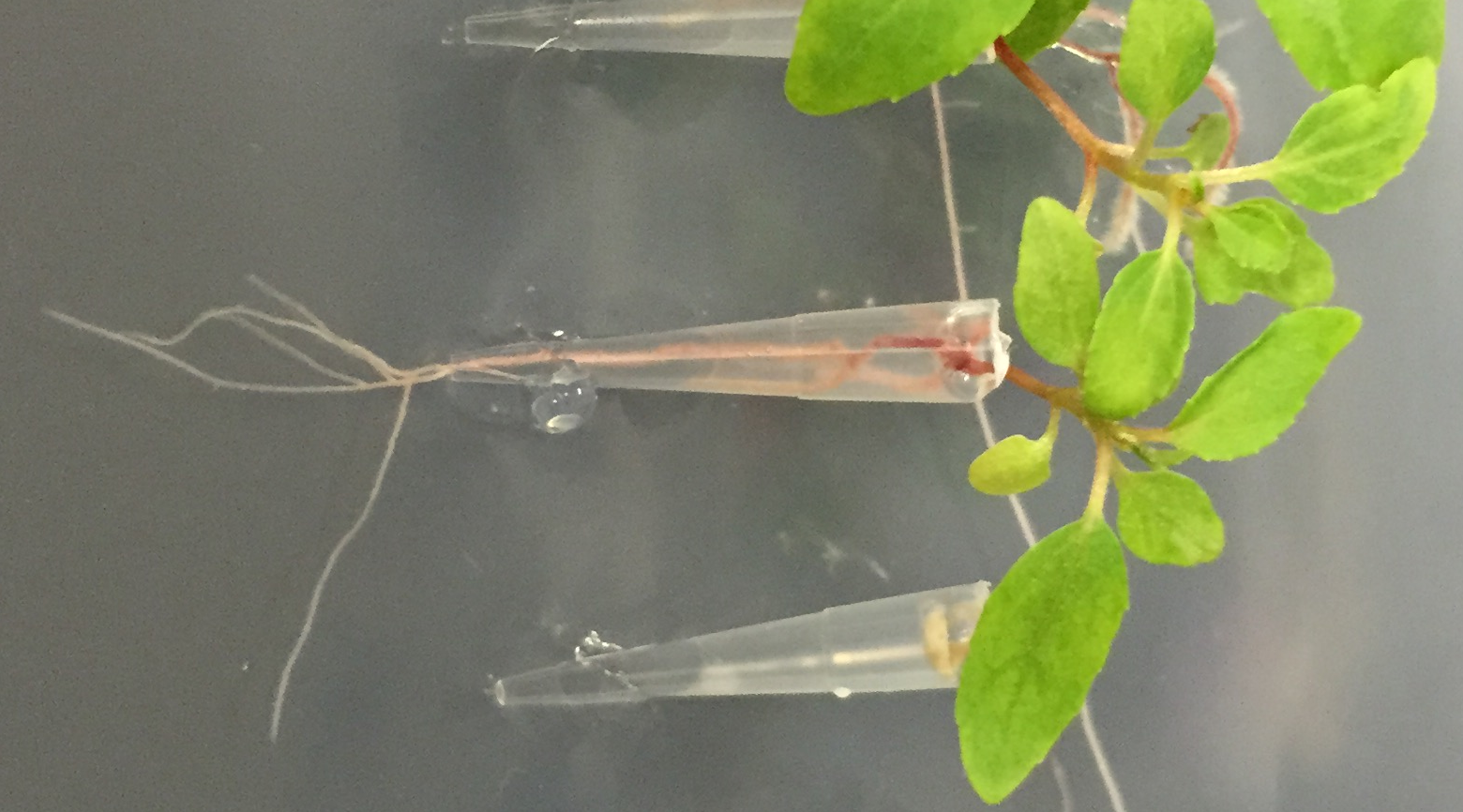

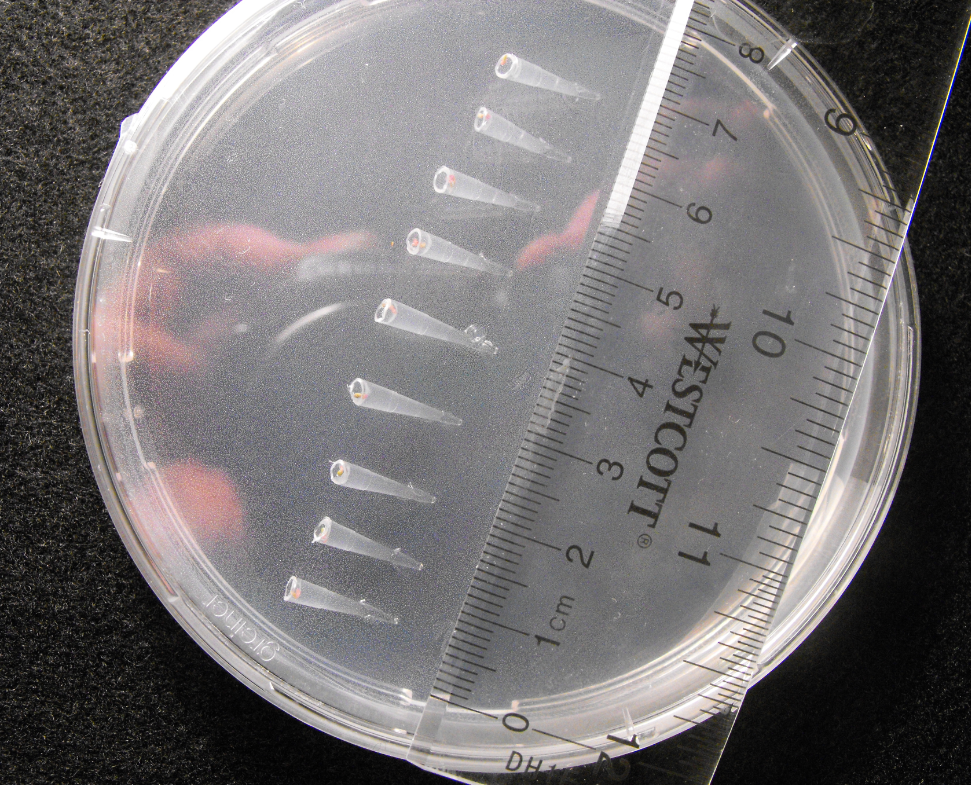
**Figure S1. Aspen seedling cultivation in pipette tips.** (a) Aspen seeds were germinated and seedlings with < 5 mm hypocotyls were transferred into 200 µL pipette tips filled with 1% Johnson’s agar. Tips were inserted at a 45° angle in a Johnson’s agar plate for gravitropic root growth and incubated in a growth chamber. (b) Healthy plant growth was observed in the pipette tips and upon exit, the primary root quickly developed branch roots (center seedling). Tips selected for mounting into the RMI-chip had a primary root reaching the end of the pipette tip (top and bottom seedlings).

b

a

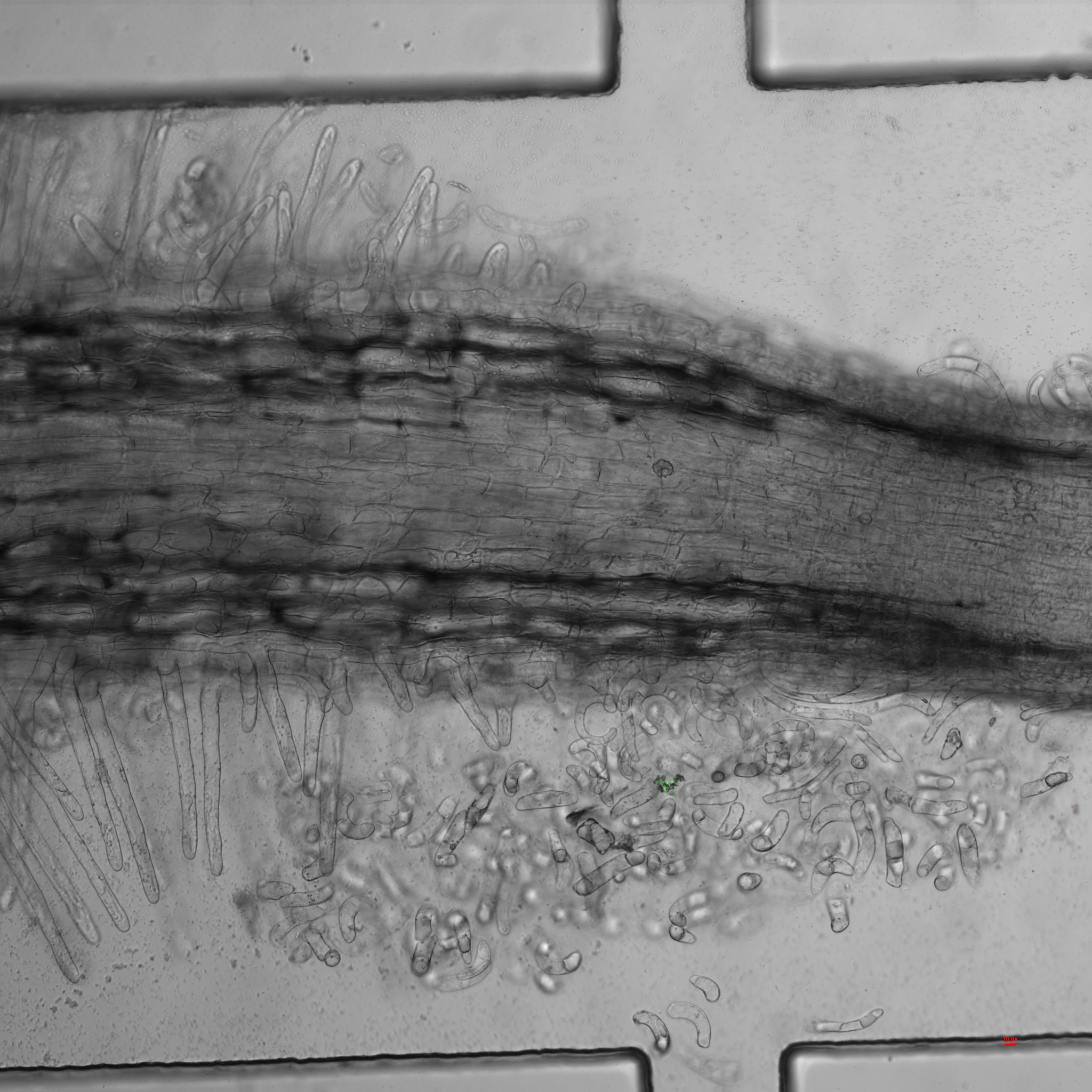

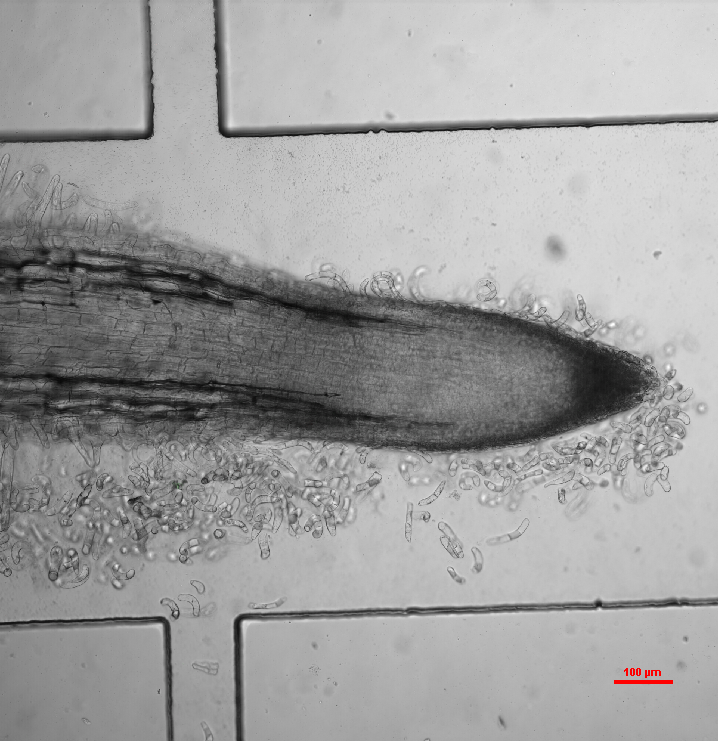
**Figure S2. Aspen seedling primary root entering the RMI-chip channel.** (a) Normal root morphology was observed in the 200 µL pipette tips. Regular primary root development was followed by branch root formation when maintained in the Petri dish with Johnson’s 1% agar. (b) Normal growth of the primary root was observed into a microfluidic channel of 80 µm x 80 µm where the root occupies all the channel space (right). (c) primary root growing semi-gravitropically in a 100 µm x 800 µm channel of the RMI-chip tilted at a 45° angle and submerged in Johnson’s solution. Within a week after mounting, the root tip reached the media inlets in the growth channel. Root hair, border cells and cell debris can be observed. At this stage, the RMI-Chip can be positioned horizontally and the flow of nutrient solution initiated.

b

200 µm

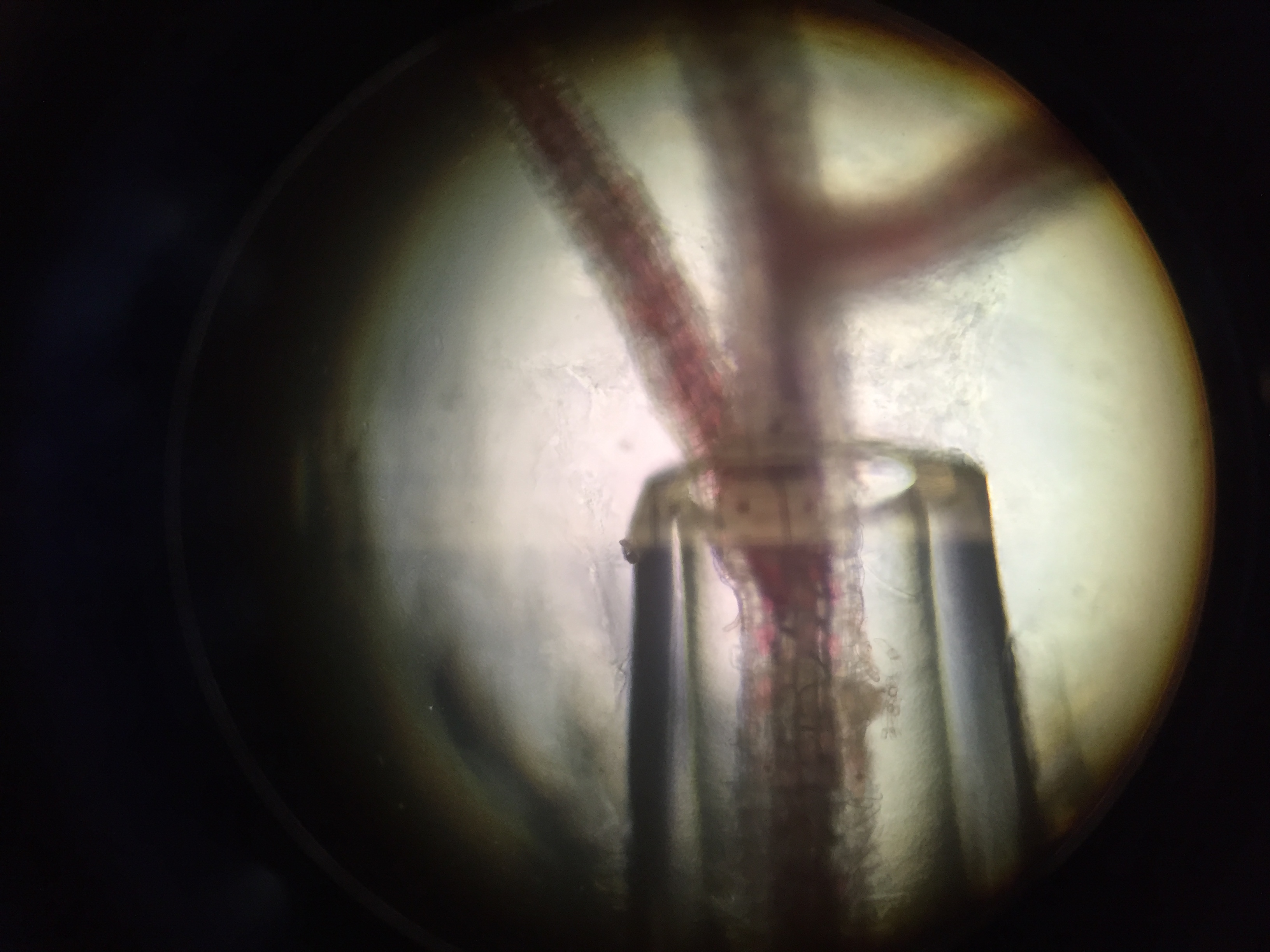

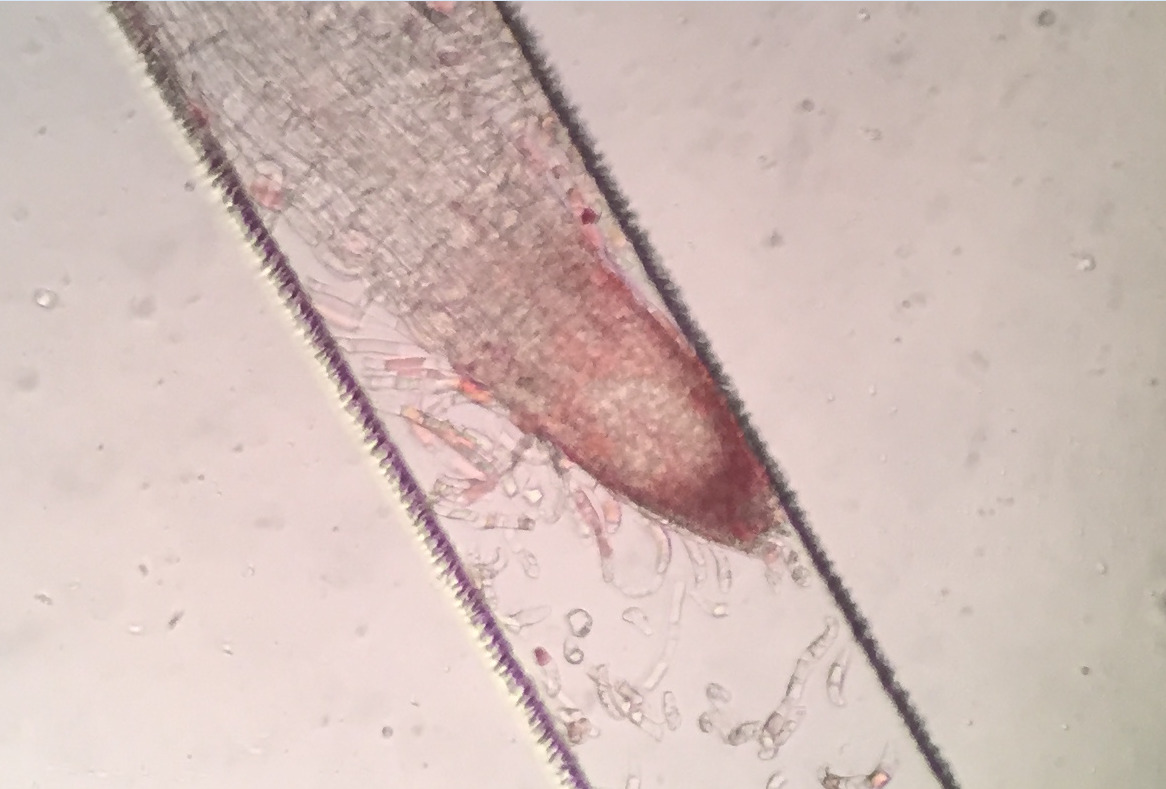

80 µm

c

a

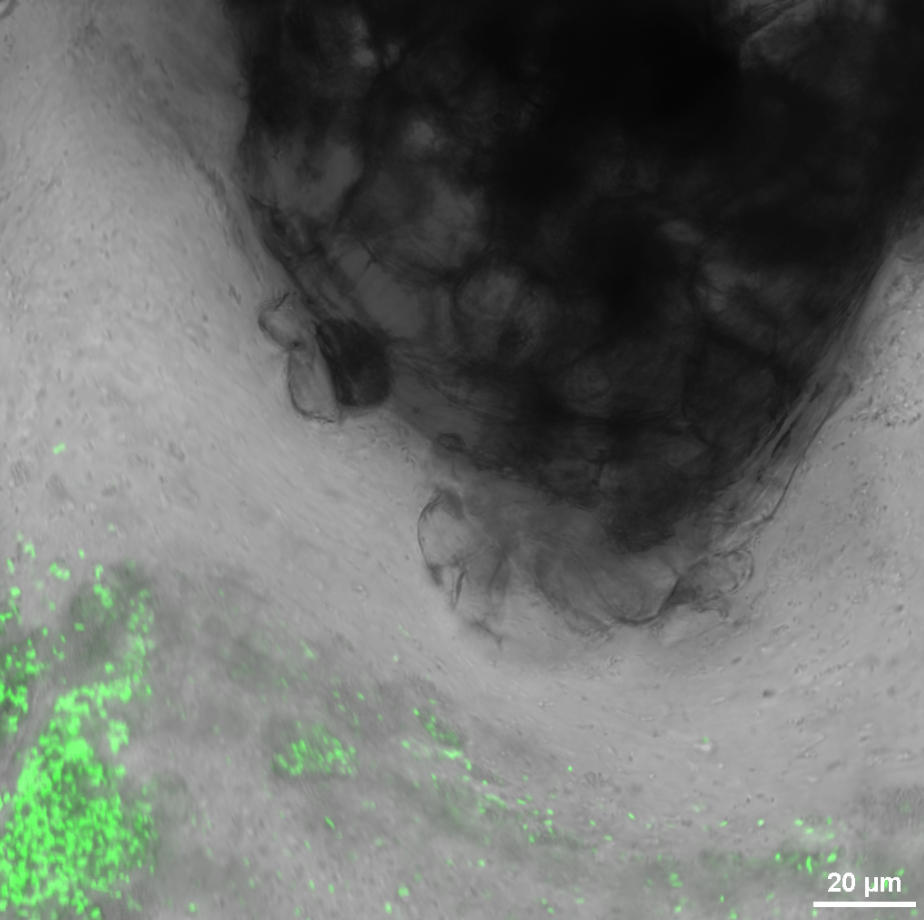

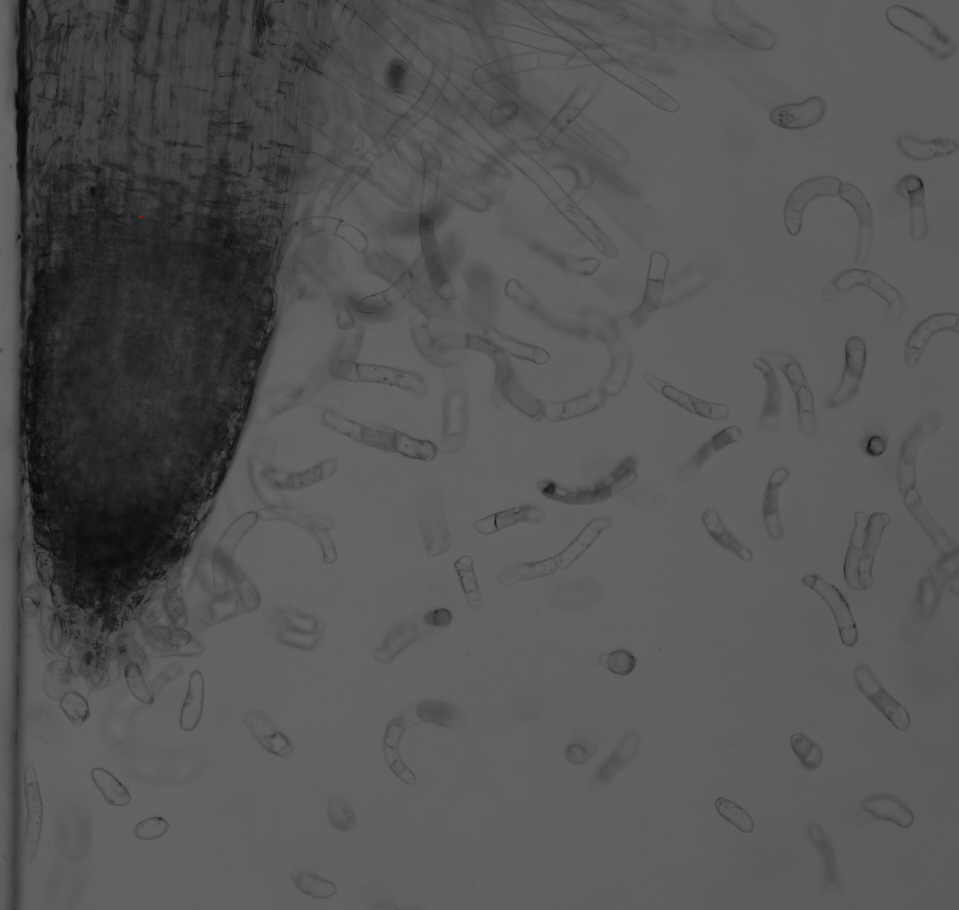

b

a

**Figure S3. Tips of a growing aspen root in absence of flow.** (a) Mucilage formed in a closed channel prevents the direct interaction of aspen seedling root tip with mNeonGreen-labeled *P. fluorescens* SBW25. (b) Aspen seedling root border cells are observed in the RMI-Chip without perfusion.

**Figure S4. RMI-Chip imaging and humidity chambers.** (a) The humidity chamber is constructed from black PMMA side panels via laser cutting (left). A sliding side window enables the removal of the imaging chamber with tubing attached. The imaging chamber has a bottom opening for direct high-resolution observations (right). (b) The modular setup with sliding windows enables continuous perfusion via a six-channel microfluidic pump connected to the RMI-Chip with FEP tubing. The imaging module is transferred with the chip during the imaging experiment.

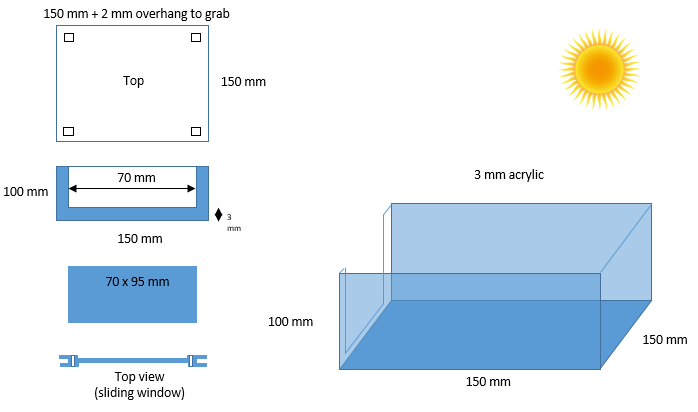

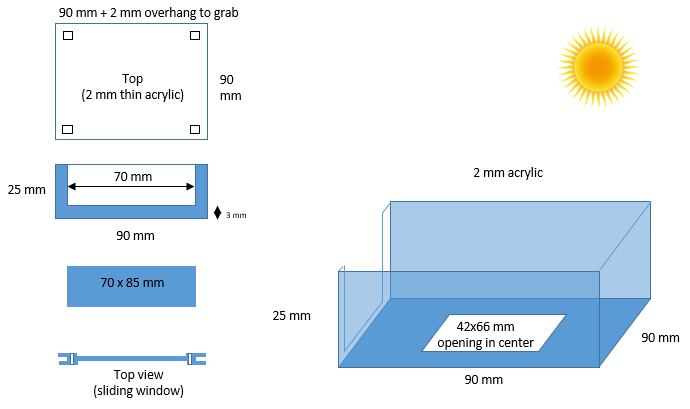

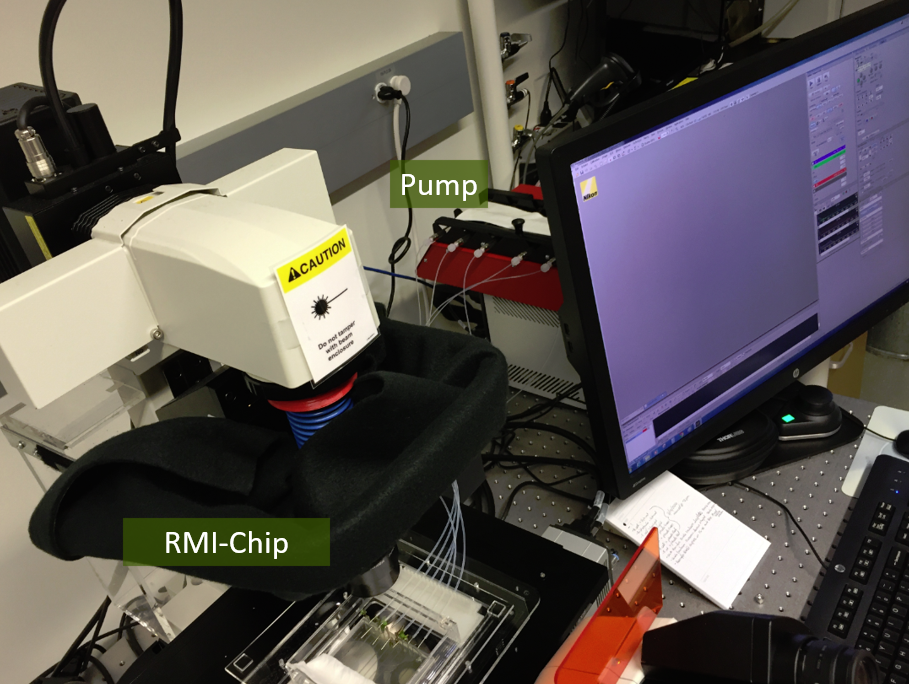

a

b

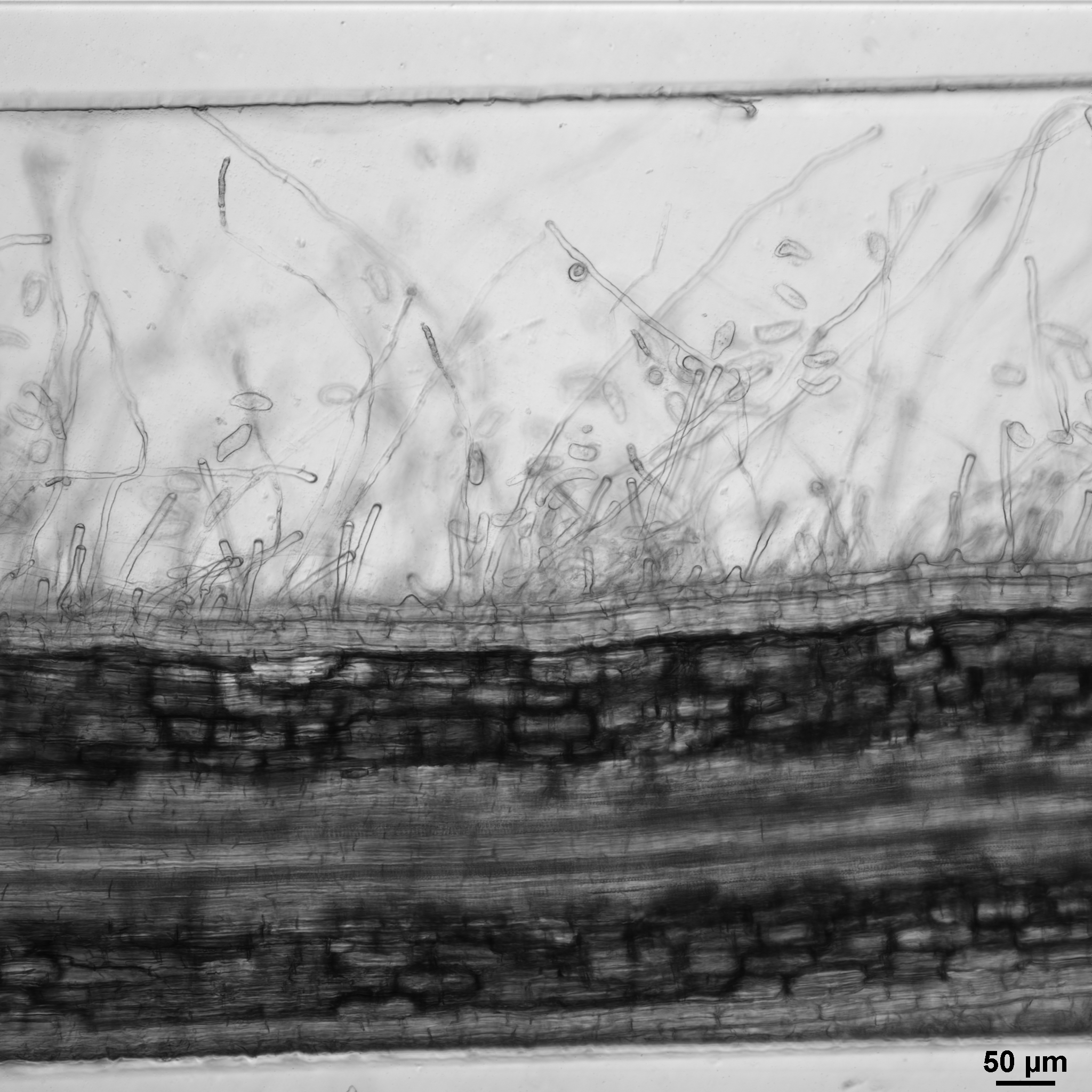

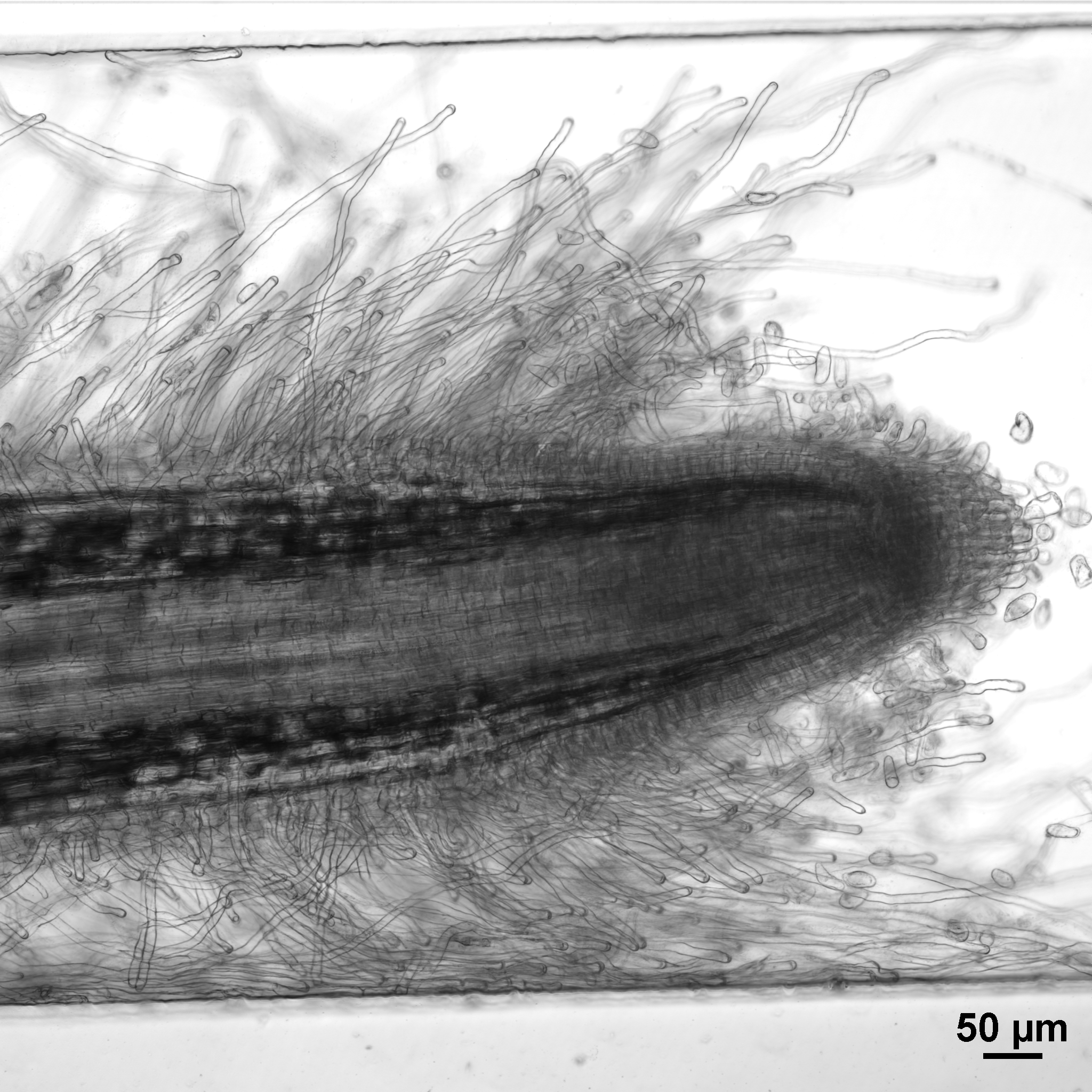

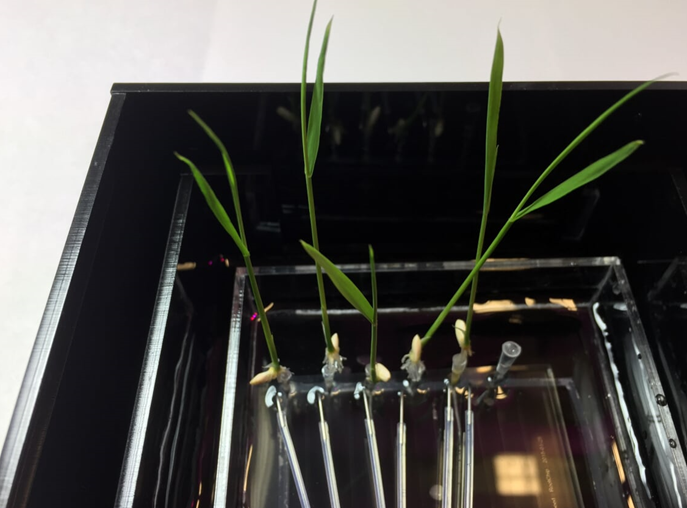

c

b

a

**Figure S5. Rice seedling in the RMI-chip.** (a) Rice seedlings grew well in the RMI-chip enabling high-resolution observations of the root. (b) The 3-chamber design is shown for long-term cultivation and repeated imaging experiments under constant perfusion with Johnson’s media. (c) Growth of rice primary root in the RMI-chip channel.

**Figure S6. Inoculation of fluorescently labeled *P. fluorescens* in the RMI-chip** (a) SBW25 cells constitutively expressing mNeonGreen were injected through the inlet and incubated 16 hours without media flow. The bacterial population appears well dispersed with some concentration of cells at the base of root hairs. (b) After continuous media flow was established (see experimental procedures), most of the bacterial cells were removed. After one day of flow, only a small number of SBW25 cells remained associated with the lower part of the primary root.

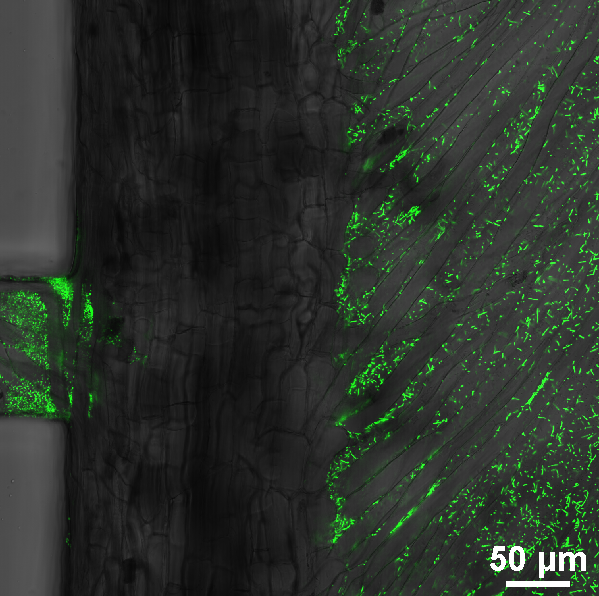

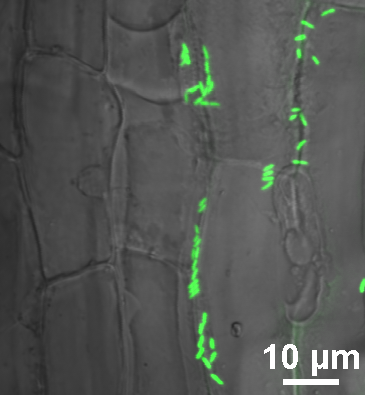

b

a

**Figure S7. Colonization of aspen primary root by *P. fluorescens* SBW25.** Five days after inoculation, mNeonGreen-labelled SBW25 cells were actively dividing and colonizing the intercellular spaces between root epidermal cells of the root cortex, and to a lesser extent, the root surface. Projection of a reconstructed 3D volume with side views show preferential occupation of deep crevices on the root surface.

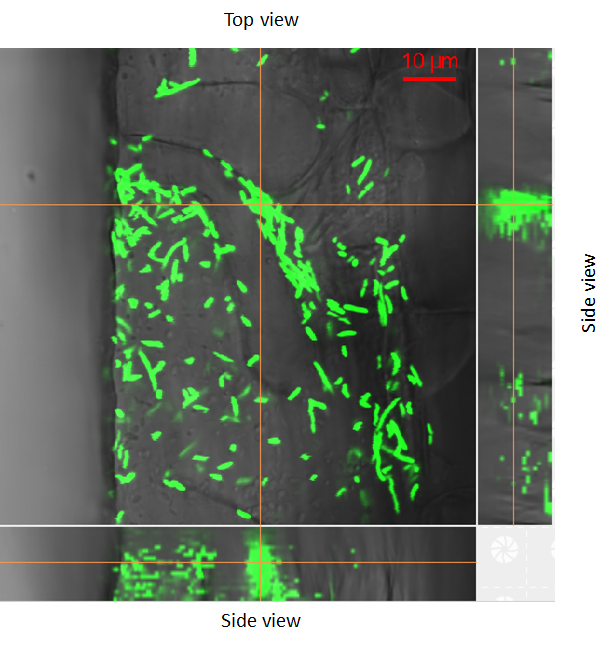

**Figure S8. Cell assemblies formed by SBW25.** (a) The inoculum including a mixture of mNeonGreen- and dsRed-labelled *P. fluorescens* SBW25 is mostly composed of elongated motile cells. (b) One day after inoculation and under flow, the root-attached cells become less elongated. (c) 13 days after inoculation, colonies of the labeled cells were observed, some with round shape microbes, resulting from tight packing. (d) Mixture of patches of microbes and loose assemblies are also observed in some regions. (e, f) Individual cells also formed quasi regularly spaced assemblies, irrespective of the label. The 3D confocal imaging revealed that the apparent coccoid shape in contrast with the normal rod-like shape of individual SBW25 cells resulted from the top view of vertically aligned cells.

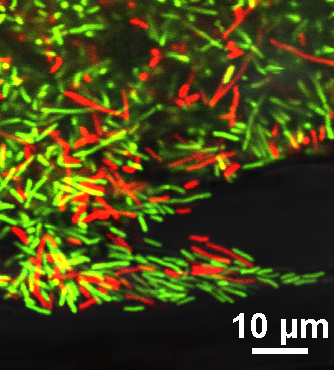

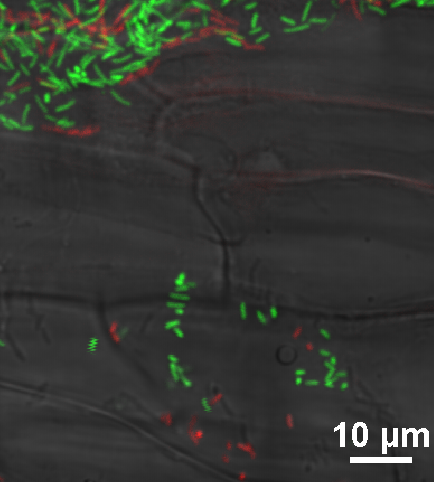

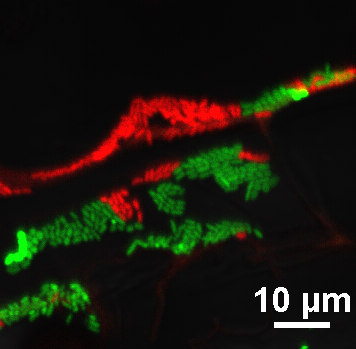

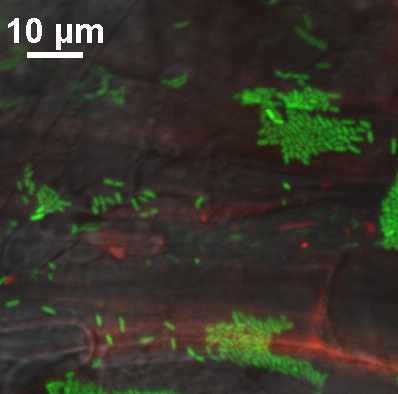

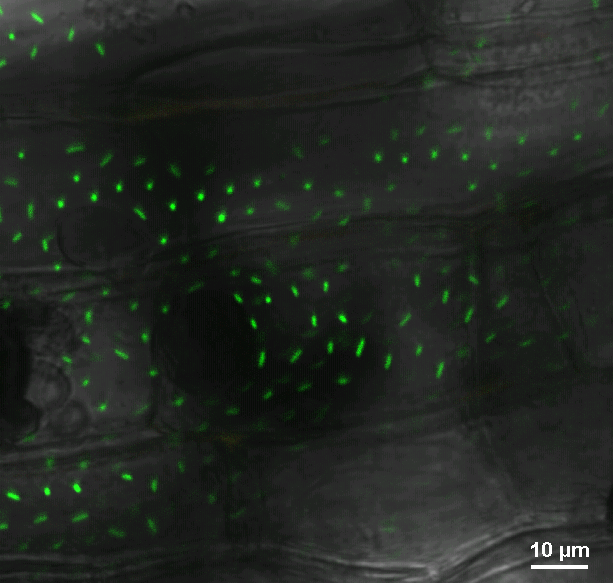

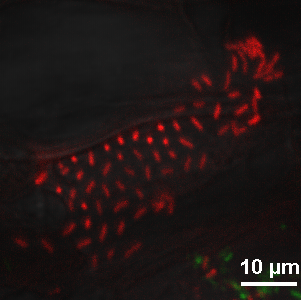

f

e

b

a

c

d

**Figure S9. Long-term root-microbe interaction.** Fluorescent SBW25 cells become rare on the aspen primary root surface 30 days after inoculation of the RMI-Chip. SBW25 cells preferentially occupy the space between root epidermal cells.

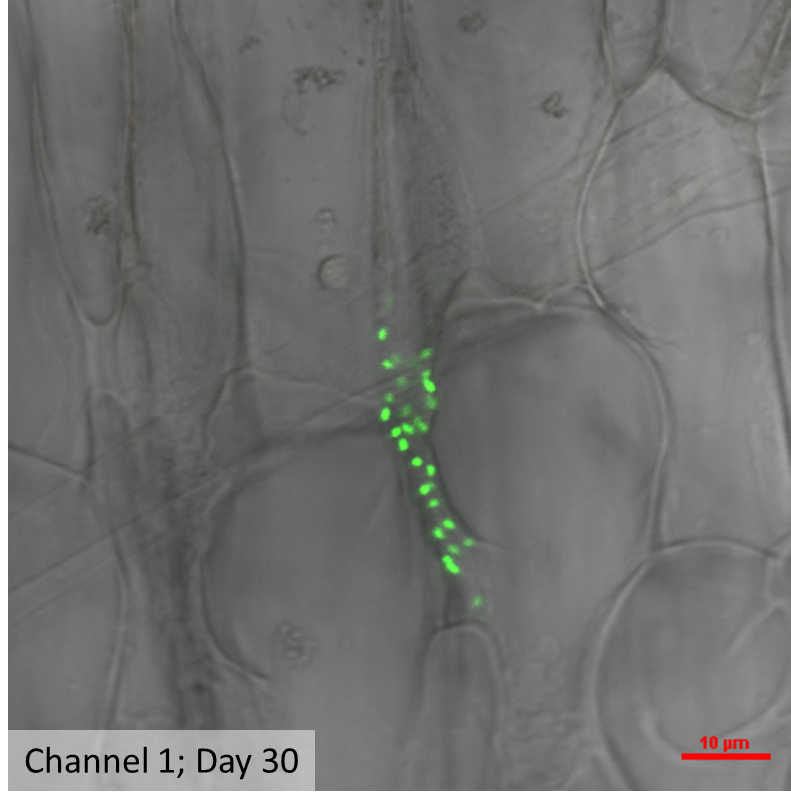
